# Evolution of a central dopamine circuit underlies adaptation of light-evoked sensorimotor response in the blind cavefish, *Astyanax mexicanus*

**DOI:** 10.1101/2024.07.25.605141

**Authors:** RA Kozol, A Canavan, B Tolentino, AC Keene, JE Kowalko, ER Duboué

## Abstract

Adaptive behaviors emerge in novel environments through functional changes in neural circuits. While relationships between circuit function and behavior have been well studied, how evolution shapes those circuits and leads to behavioral adpation is poorly understood. The Mexican cavefish*, Astyanax mexicanus*, provides a unique genetically amendable model system, equipped with above ground eyed surface fish and multiple evolutionarily divergent populations of blind cavefish that have evolved in complete darkness. These differences in environment and vision provide an opprotunity to examine how a neural circuit is functionally influenced by the presence of light. Here, we examine differences in the detection, and behavioral response induced by non visual light reception. Both populations exhibit photokinetic behavior, with surface fish becoming hyperactive following sudden darkness and cavefish becoming hyperactive following sudden illumination. To define these photokinetic neural circuits, we integrated whole brain functional imaging with our *Astyanax* brain atlas for surface and cavefish responding to light changes. We identified the caudal posterior tuberculum as the central modulator for both light or dark stimulated photokinesis. To unconver how spatiotemporal neuronal activity differed between surface fish and cavefish, we used stable pan-neuronal GCaMP *Astyanax* transgenics to show that a subpopulation of darkness sensitve neurons in surface fish are now light senstive in cavefish. Further functional analysis revealed that this integrative switch is dependent on dopmane signaling, suggesting a key role for dopamine and a highly conserved dopamine circuit in modulating the evolution of a circuit driving an essential behavior. Together, these data shed light into how neural circuits evolved to adapte to novel settings, and reveal the power of *Astyanax* as a model to elucidate mechanistic ingiths underlying sensory adaptation.

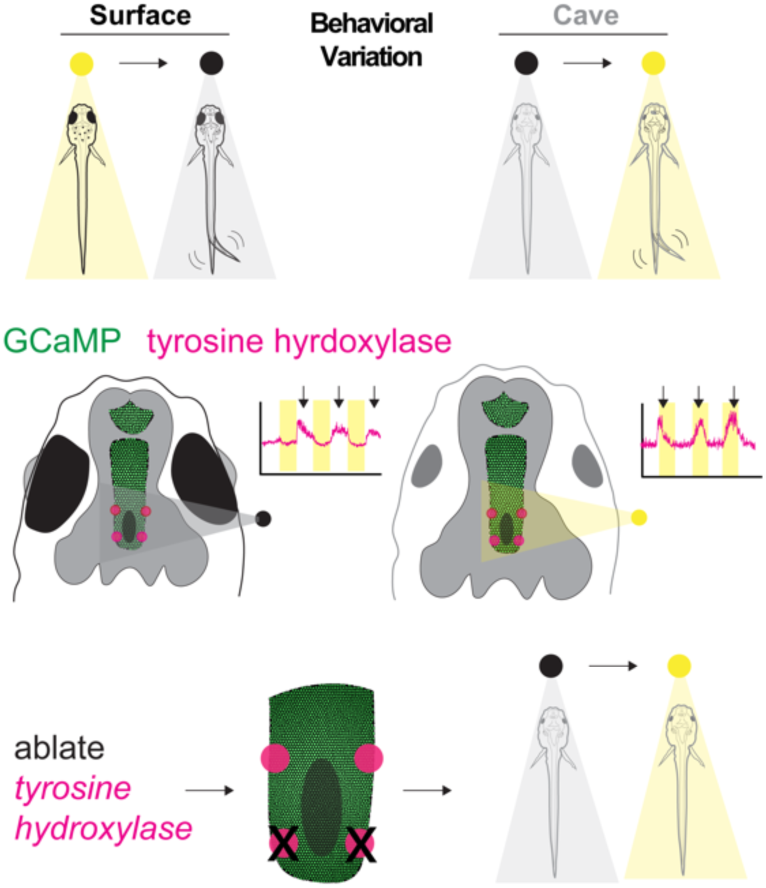

## Introduction

Behavioral responses are driven by the dynamic coordination of brain-wide neural circuits, which act in coordinated fashion to take in sensory information and elicit an appropriate behavioral response. Behaviors are finely tuned to the environment that an organism lives in, and neural circuits underlying these responses are under strict evolutionary pressure [1–4]. Model systems have significantly enhanced our understanding of how neural circuits develop and modulate behavior [5–7]. Studies using circuit ablations or optogenetics in wide ranging systems from flies and fish to mammals have revealed the effect of dysregulated neural circuits and their impact on behavior. However, less is known about how the circuits evolve to give rise to adaptative behaviors [4, 8, 9]. While advances in genetic technology in model systems has made it possible to inactivate or over activate circuits, a significant impediment to understanding how circuits lay in the inability to modify neural circuits to change a behavior. Our understanding of how evolution shapes circuits comes largely from comparing neural circuit function across closely related species that inhabit similar environments, though large divergence times and incomplete evolutionary histories makes inferring generalizable models of how neural circuits evolve difficult [7, 10–16]. Addressing this question requires a species with evolutionarily divergent forms with complete evolutionary histories, robust behavioral differences, accessibility to brain-wide neural circuits, and amenability to genetic and neuronal interrogation [4].

The blind Mexican tetra (*Astyanax mexicanus*) is an emerging model for understanding the evolution of neural circuits and behavior. *A. mexicanus* exists as surface dwelling river fish and more than 30 independently evolved cave populations, which have adapted to hundreds of thousands of years of nutrient scarcity and perpetual darkness [17–20]. While surface populations have retained eyes and pigment, cave populations share eye degeneration, loss of pigment and exhibit variation in sleep, stress, feeding and metabolism [21–25]. These evolved traits can be functionally investigated genetically by interbreeding surface and cave morphs, producing viable hybrid offspring that permits investigation of the genetic basis of trait variation, including genetic mapping [26–30]. Recently, we have established transgenesis and targeted genome mutagenesis in surface and cavefish populations. This work has brought Astyanax into the genetic era, and has resulted in a number of stable lines such as ones that express the Ca2+ sensor GCaMP pan-neuronally [31]. This specific transgenic line allows for functional imaging of neural response to sensory stimuli. Together, this model is uniquely poised to understand how evolution impacts circuit function and behavioral adaptation.

Sensory systems provides a platform for understanding the evolution of behaviorally relevant traits in response to a changing environment [32]. In zebrafish, detailed work examining the behavioral and neuronal response to changes in illumination has been particularly well studied. When fish in an illuminated background are exposed suddenly to darkness, fish become hyperactive and engage in a light searching behavior[11, 33–36] [37]. This behavior is presumed to be an evolutionary adaptation which allows visual perception to guide fish’s behavior. Moreover, recent work has shown that such photokinesis is evolutionarily conserved in sighted fish spanning more than 200 million years of evolution[38, 39]. While photokinesis has not been examined in Astyanax, previous work has shown that non-visual detection of changes in illumination is preserved in both surface and cave Astyanax.

In this study, we investigated how evolution shapes the sensorimotor processing of light stimuli in the blind Mexican cavefish (*Astyanax mexicanus*) by comparing photokinetic behavior and neural circuits across surface fish and cavefish populations. We show that cavefish develop a light photokinetic response, becoming active in the light, in contrast to the ancestral dark photokinetic response observed in surface fish, becoming active in the dark, which represents a fundamental sensorimotor adaptation to evolving in the perpetual darkness of a cave. Using whole-brain functional mapping of neural activity with stable GCaMP surface and cavefish, we identified a neural circuit involving the hypothalamus that drives cave-specific differences in sensorimotor integration and show the emergence of a novel functional cell type in this region. Finally, we show that a set of dopaminergic neurons, which are conserved from fish to humans, exhibit responses to light cues that functionally impact the photokinesis. This study reveals that the integration of light stimuli can be transformed through the process of evolution, producing a behavioral adaption via changes in the functional properties of a deeply conserved dopaminergic cell type.

## Results

### Identification of a fundamental sensorimotor in cavefish

Zebrafish and other sighted teleost exposed to sudden periods of darkness become hyperactive and display a ‘light searching’ behavior [37, 40, 41]. To test these responses in surface and cave *Astyanax*, we devised a similar assay to record locomotor behavior under alternating periods of 5-min lights on followed by 5-min lights off (Figure 1a). Surface fish showed baseline locomotor patterns under periods of illumination, and hyperactivity during dark periods. Conversely, we found that cavefish had baseline locomotion in darkness and hyperactivity when the lights were turned on (Figure 1b), suggesting a change in their preference to illumination. To quantify how strongly each fish responded to lights on or lights off, we devised a metric, the photokinesis index, by dividing the difference of light on activity (light off - light on) by the sum of light off activity (light on + light off), called (Figure 1a, see methods). A positive photokinesis index indicates the fish was hyperactive when the lights were turned off, whereas a negative photokinesis index suggests fish became hyperactive when lights were turned on. We found that both zebrafish (*Danio rerio*) and *Astyanax* surface fish exhibit positive photokinesis indices consistent with their hyperactivity during the lights off phase, (Figure 1c, Supplementary Figure S1, Supplementary Table S1&2), whereas cavefish exhibit negative photokinesis indices, suggesting they are hyporeactive during the lights on phase (Figure 1b&c, Supplementary Figure S1, Supplementary Table S1-3). These results reveal that evolution in perpetual darkness alters photokinetic responses in behaviorally relevant ways.

**Figure 1.**
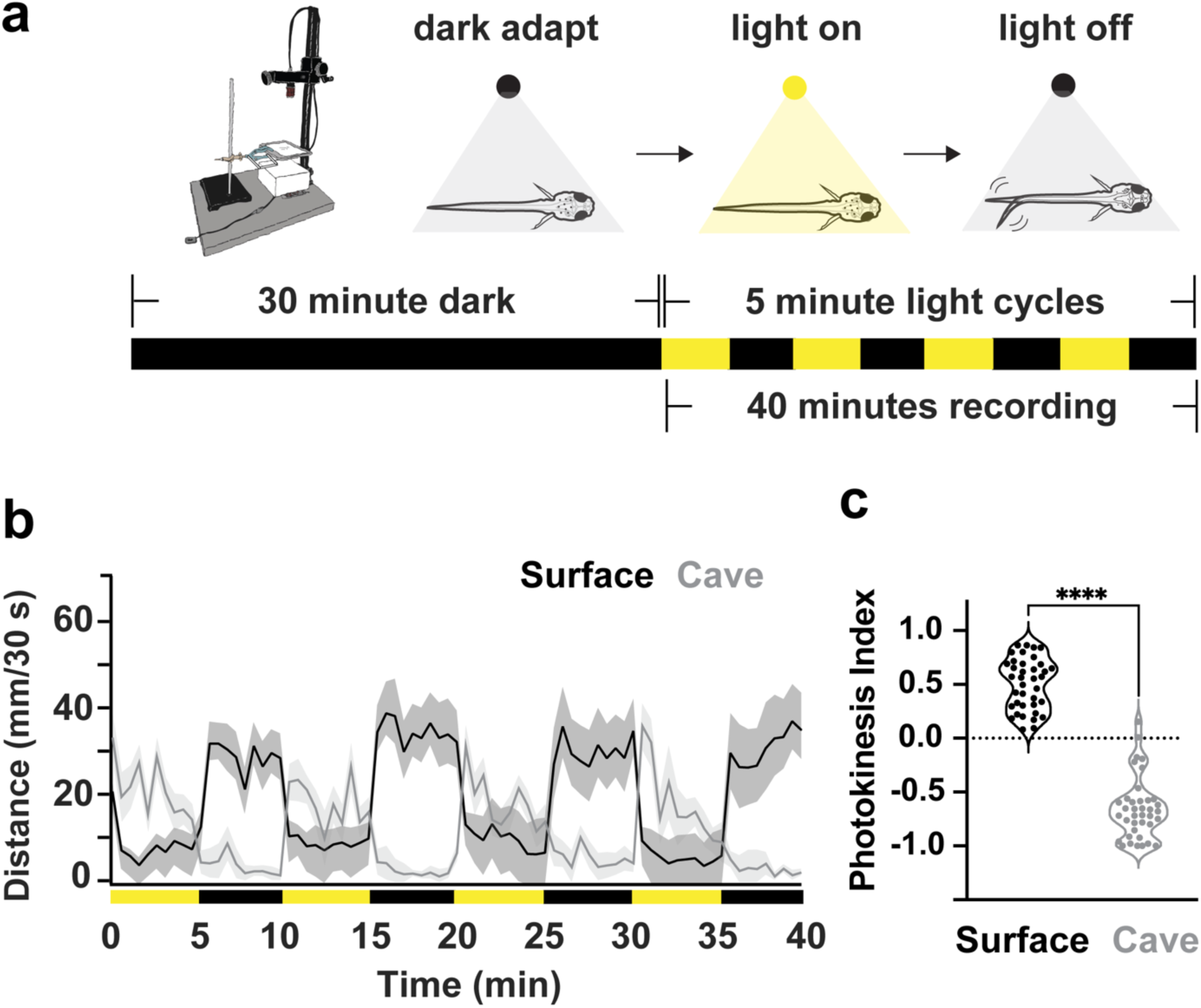
Cavefish exhibit a negative photokinesis index in comparison to a positive photokinesis index in surface fish. **a** Diagram illustrating eyed fish behavior and the light cycles used in the photokinesis experiments. Surface fish exhibit hyperactive behavior in dark cycles. Larvae were dark adapted before 5 minute alternating cycles of light on (yellow) to light off (black). **b** Line graph showing the inverse behavioral patterns of hyperactive behavior for cavefish (lights on=yellow bars) and surface fish (lights-off=black bars). **c** Photokinesis index box plots quantifying relationship between increased activity and light cycle (positive = light off, negative = light on). **d.** Average velocity of 30 second binned light on to light off transitions. Sample sizes, surface fish (n=42) and cavefish (n=34). Statistical significance represents output from a Tukey multiple comparisons corrected one-way ANOVA. ** = p<0.01, **** = p<0.0001.

### Whole brain functional mapping reveals differences in forebrain neural activity underlies light preference

A functional sensory circuit typically contains neurons to receive input, neurons that process that input, and neurons to produce output [16, 42]. We sought to identify the neurons that are active during light and dark periods in both cave and surface fish forms. To characterize this sensorimotor circuit in surface fish and cavefish, we performed phospho-ERK (pERK) MAP-mapping of the entire *Astyanax* brain [43–45], comparing neural activity between lights-on and lights-off transitions within surface fish and cavefish populations. pERK is used as a proxy for neural activity, exhibiting peak phosphorylation within 3-5 minutes of neuronal firing [43]. Groups of fish were split into two groups, exposed to either lights-off or lights-on for 5 minutes, stained immunohistochemically for tERK and pERK and then compared within a population (e.g., surface fish or cavefish). For light receptive regions, we found increased neural activity in the optic tectum of surface fish when the light was turned on, while the pineal body showed increased neural activity when the lights were turned off, recapitulating photokinesis activity maps in zebrafish [46] (Figure2a&b). Alternatively, cavefish did not exhibit increased activity in light receptive regions, but did show increased activity in the pineal body during lights-off (Figure2a&b). Due to its standing role as a light sensing organ [47, 48], the pineal is the likely anatomical locus sensing changes in illumination in both cavefish and surface fish.

We next asked what brain region could be central to deep brain light processing. We reasoned that the region processing light would show enhanced neural activity in surface fish when the lights turned off, whereas the same area would increase neural activity in cavefish when the lights turned on. We found two anatomical areas in the ventral forebrain that matched this expectation. When registered to the *Astyanax* brain atlas [45], we found that this area mapped to the preoptic region of the hypothalamus (blue symbol) and the posterior tuberculum (red symbol; Figure 2d&e). Together, this suggests a neural circuit involving the pineal as a light sensing organ, and the hypothalamus and posterior tuberculum as processing centers. Moreover, that there were no significant differences in pERK staining of the pineal, but rather opposite activity profiles for the central sensory processor suggest that environmental pressure is acting on the central processor of light and not a primary light sensing organ.

**Figure 2.**
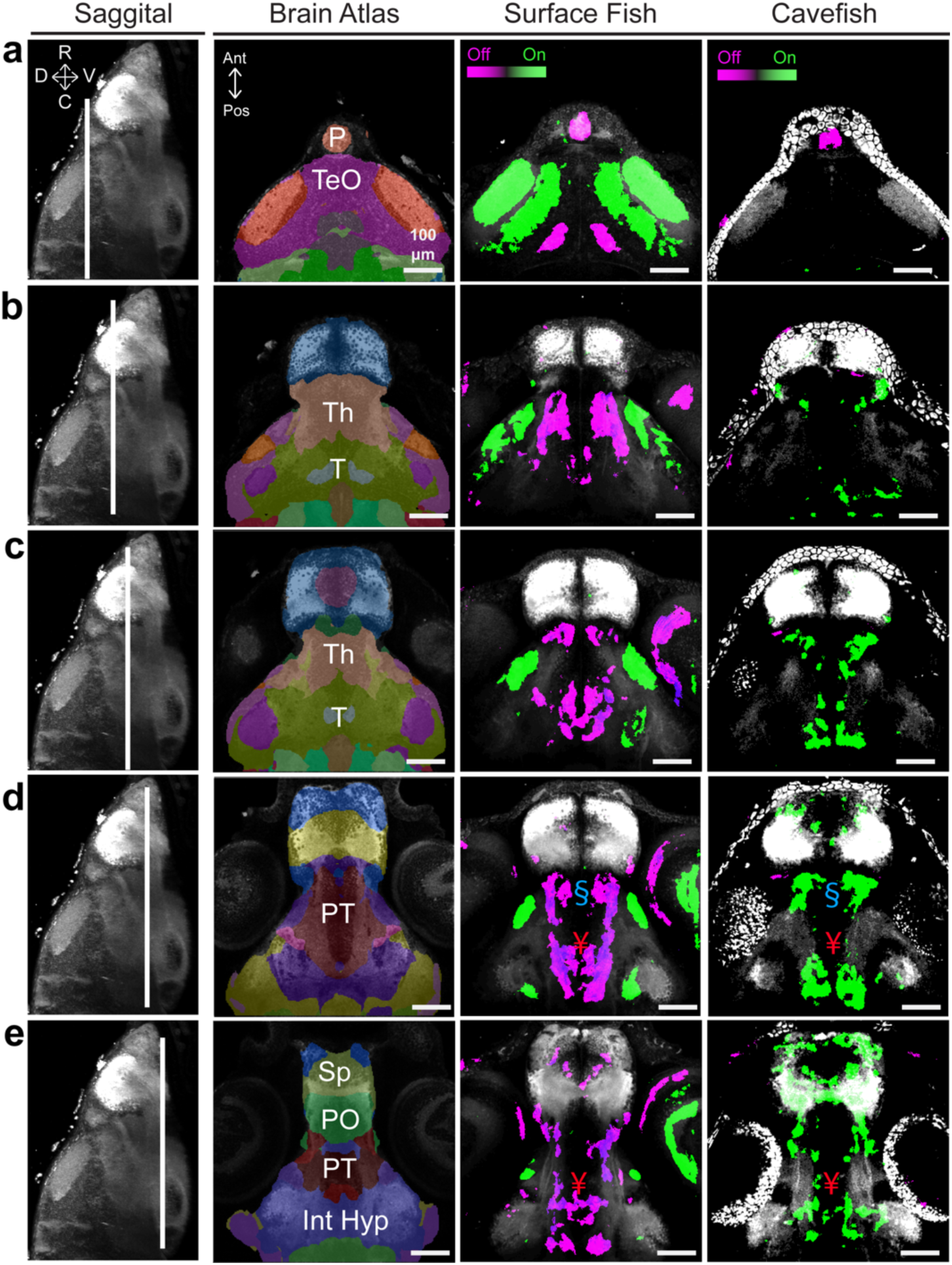
Brain mapping identifies overlap and variation of neural activity in the ventral forebrain between surface fish and cavefish. The first column displays a sagittal maximum projection through the middle of the brain, with directional arrows representing dorsal (D), ventral (V), rostral (R) and caudal (C). The second column provides an optical section through the *Astyanax* brain atlas and tERK reference brain. The last two columns contain pERK neural activity maps displaying regions of increased pERK immunostaining during lights off (magenta) or lights on (green) in surface fish and cavefish. Rows represent dorsal to ventral optical sections through the (**a)** pineal (P) and optic tectum (TeO), (**b&c)** thalamus (Th) and tegmentum (T), and finally (**d&e)** the subpallium (SP), preoptic region (PO), posterior tuberculum (PT) and intermediate hypothalamus (Int Hyp). Sample sizes, surface fish (n=18) and cavefish (n=20).

### Analysis of hybrid fish suggests photokinesis is a genetically heritable trait

A strength of the Astyanax system is the ability to cross different populations to one another, and study the genetics that underly complex traits. We first crossed a pure surface fish with a cavefish isolated from the Pachón cave to produce a brood of F1 hybrid fish. Compared to pure surface and cavefish, F_1_ exhibited an intermediate photokinetic phenotype (Supplemental Figure S2, brown circles). We then crossed F1 fish with one another to produce a population of F2 hybrid fish, and subjected these larvae to our photokinesis assay. We found varying levels of photokinesis in F2 hybrids with some progeny reacting more similarly to surface fish, and others reacted similarly to cavefish (Figure 3b&c, F_2_ hybrids are blue circles). When examined as a population, F2 hybrid fish spanned the entire range of surface to cave. The photokinetic mean of F_2_ hybrid fish was near zero (−0.0355), with a much higher variability than either pure breeding population (S=0.3754, C=-0.3687, F_2_ = −0.0355), measure consistent with a genetically encoded trait. These results suggest photokinesis differences are driven by complex, multi-locus allelic variation between wild surface fish and cavefish populations.

**Figure 3.**
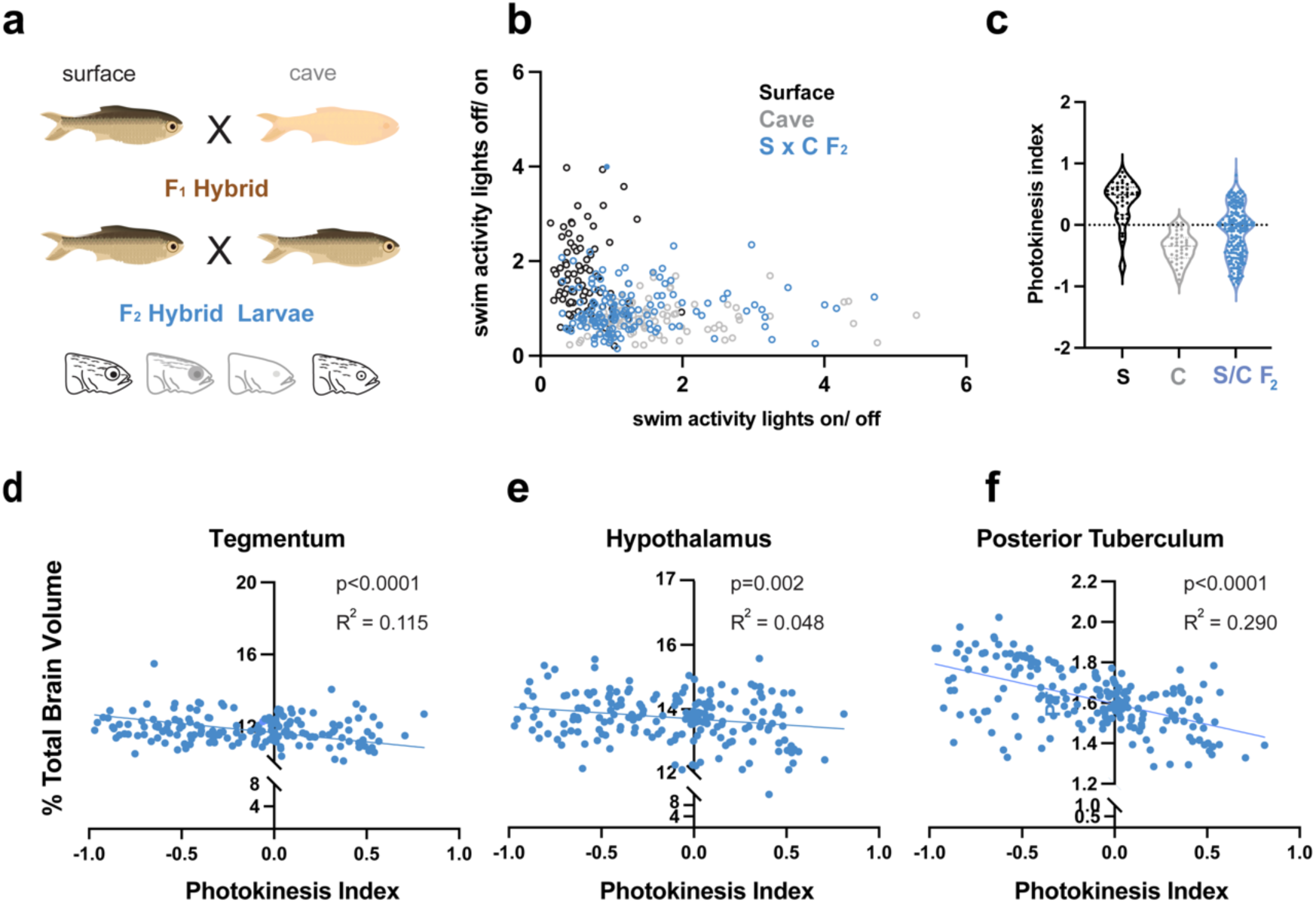
Photokinesis behavior in F_2_ hybrids is highly variable and photokinesis indices are negatively correlated to brain region size. **a** Diagram illustrating the crossing scheme for producing F_2_ hybrid larvae to correlate behavior to biological traits. **b** Scatter plot displaying the relationship between light-on and light-off transition states. Surface fish are grouped along the y-axis, cavefish along the x-axis, and F_2_ hybrids are scattered across both grandparental populations. **c** Photokinesis index violin plots exhibiting the range of behaviors of F_2_ hybrids, from positive to negative photokinesis values. Scatter plot of brain region volume against photokinesis behavior for (**d**) tegmentum, (**e**) hypothalamus and (**f**) posterior tuberculum Sample sizes, surface fish (n=42), cavefish (n=34), surface to cave F_2_ hybrids (n=199).

### Brain-wide analysis of neuroanatomy in hybrids reveals correlation between photokinesis behavior and posterior tuberculum size

Due to the established relationship between functional variation and anatomical differences within a population [49, 50], we next asked whether changes in neuroanatomical volume were driving the differences in photokinesis. To determine whether brain region volume drove differences between surface fish and cavefish photokinesis behavior, we assessed light-evoked activity in F_2_ hybrid fish, and then examined correlations with this behavior with the volume of every known neuroanatomical region using a recently published Astyanax brain atlas[45]. Each individual hybrid larva was behaviorally tested and scored in our photokinesis assay, stained for brain-wide neuroanatomy using an ERK antibody, and computationally analyzed for regional brain volume. We then performed a parametric correlation analysis between neuroanatomical volume and photokinesis indices (Figure 3, Supplementary Figure S3). Surface-cave F_2_ larvae exhibited negative correlations between photokinesis indices and volume of the tegmentum, hypothalamus and posterior tuberculum (Figure 3d-e, Supplementary Tables S-6), supporting a role for anatomical evolution of the ventral forebrain in modulating these behaviors. In contrast, no other brain region showed a significant correlation between photokinesis indices and regional volume (Supplementary Figure S3, Supplementary Table S7-16). These results suggest light and dark photokinesis is likely influenced by anatomical variation of the ventral forebrain in *Astyanax* populations.

### Neurons in the posterior tuberculum exhibit novel off to on responses in cavefish

Functional *in vivo* imaging provides a powerful tool to examine the relationship between neural activity and behavior. We sought to determine whether spatiotemporal neural activity of cells within the posterior tuberculum are differentially modulated by light in surface and cavefish. Our lab has developed stable transgenic *Astyanax* lines that express the genetically encoded calcium indicator GCaMP6s in nearly every neuron of the brain throughout the brain (*Tg(_zfish_elavl3:H2B-GCaMP6s)*) [51, 52]. Larvae were mounted in a drop of low melt agarose, placed under a confocal microscope, and subjected to 30 second alternating periods of lights-on and lights-off (Figure 4a). We found that the posterior tuberculum contained light responsive clusters of neurons (Figure 4b&c). Importantly, each cluster in surface fish showed a light-off tuned response, and the patterns of activity among clusters was stable across different larvae (Figure 4b, black arrows in grey stripes). By contrast, while clusters of neurons in the medial portion of cavefish were also light off responsive, those clusters in the caudal portion of the posterior tuberculum exhibited light on responses, consistent with the hyperactivity in response to illumination in this morph (Figure 4c, black arrows in yellow stripes). To quantify variation across morphs, we calculated calcium transient averages for clusters 4 and 5 and compared across light conditions (Figure 4d, Supplementary Tables S17-20). Lastly, in both surface fish and cavefish, rostral regions exhibited no clear light-evoked activity (Supplementary Figure S4). Therefore, the posterior tuberculum is unique in its differential response to light between surface and cavefish populations.

**Figure 4.**
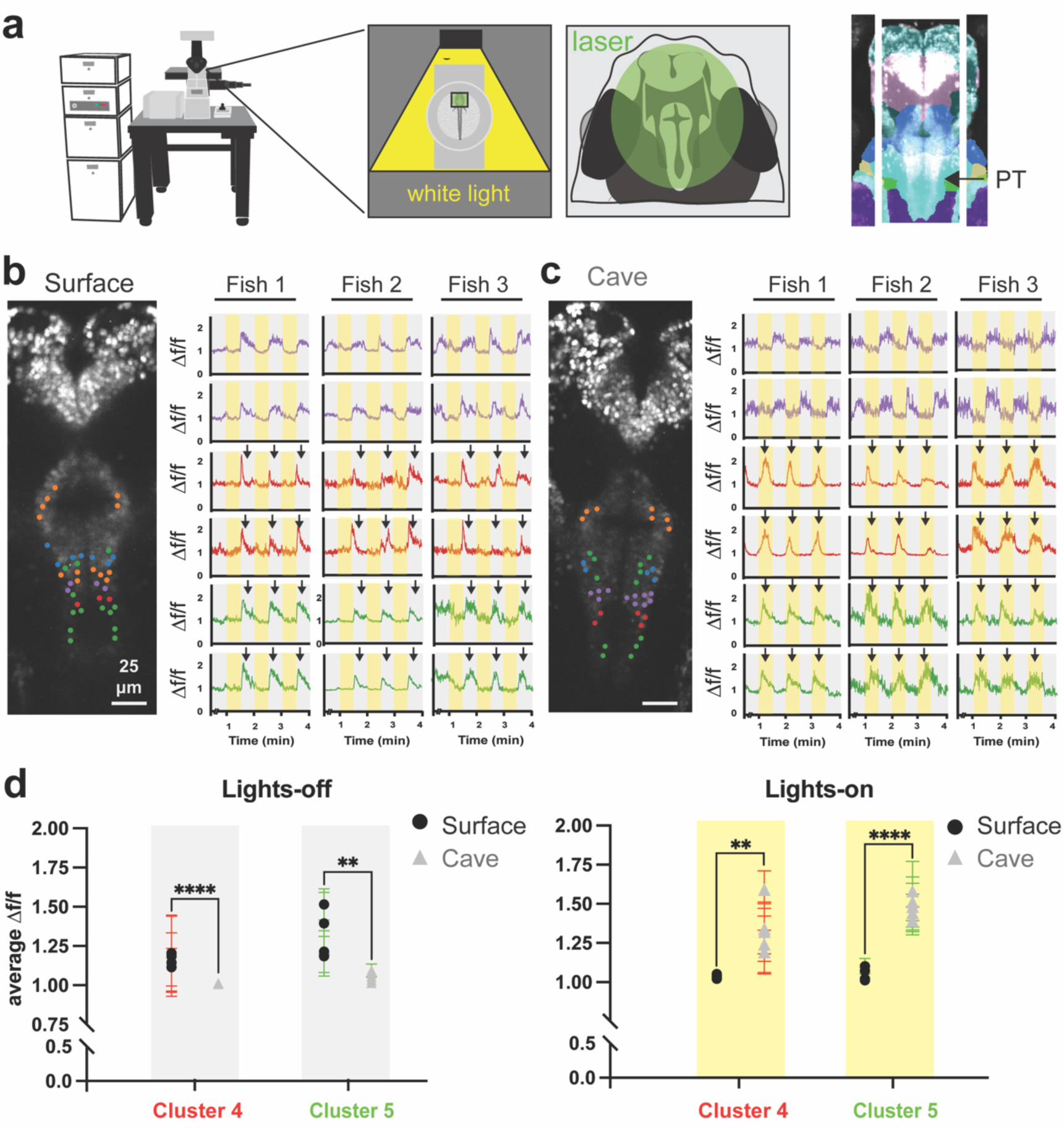
Clusters of neurons in the posterior tuberculum of cavefish exhibit light-on tuning in comparison to light-off tuning in surface fish. **a.** Diagram illustrating the experimental setup of light stimulated agarose embedded surface and cavefish GCaMP6s larvae. Projection inset shows the segmented region analyzed in the live imaging dataset. Surface fish (**b**) and cavefish (**c**) timeseries clusters based on stimulus tuning, dark cycles (grey stripes) or light cycles (yellow stripes). Light condition peaks are noted by black arrows for cluster 4 (red) and cluster 5 (green). Coronal projections represent analyzed regions and display color coded neuronal clusters. **d** Box plots comparing average delta f over f for lights-off and lights-on conditions of clusters 4 and 5 in the caudal posterior tuberculum. Sample size, surface fish (n=4) and cavefish (n=5). Statistical significance represents t-test comparisons between populations for each cluster according to light conditions, ** = p<0.01, *** = p<0.001, **** = p<0.0001

### Neurons in the caudal posterior tuberculum that are dark tuned in surface fish and light tuned in cavefish express dopaminergic markers

The posterior tuberculum is comprised of many different cell types, yet one of the most conserved are the dopaminergic neurons of the descending diencephalic system. These cells are thought to be analogous to the A11 Dopamine cells in mammals, and modulate diverse internal states including stress, mood, and reward [53, 54]. To assess whether light-evoking cells in *Astyanax* were dopaminergic, we employed a unique method that combines live GCaMP imaging with subsequent RNA *in situ* hybridization to molecularly identify neurons that are active in light or dark conditions. Briefly, following live GCaMP imaging, larvae were fixed in agarose to preserve the brains position and probed for the dopaminergic marker, *tyrosine hydroxylase* (*th*, Figure 5b). Live to fixed registration of GCaMP channels revealed that caudal neurons of the posterior tuberculum with opposite dark to light stimulus tuning express *tyrosine hydroxylase* (Figure 5c-e). These data suggest that the descending diencephalic system could be integrating light signals and sending the information to hindbrain motor areas, and this conserved set of neurons may be responsible for differences in light preference between surface and cavefish.

**Figure 5.**
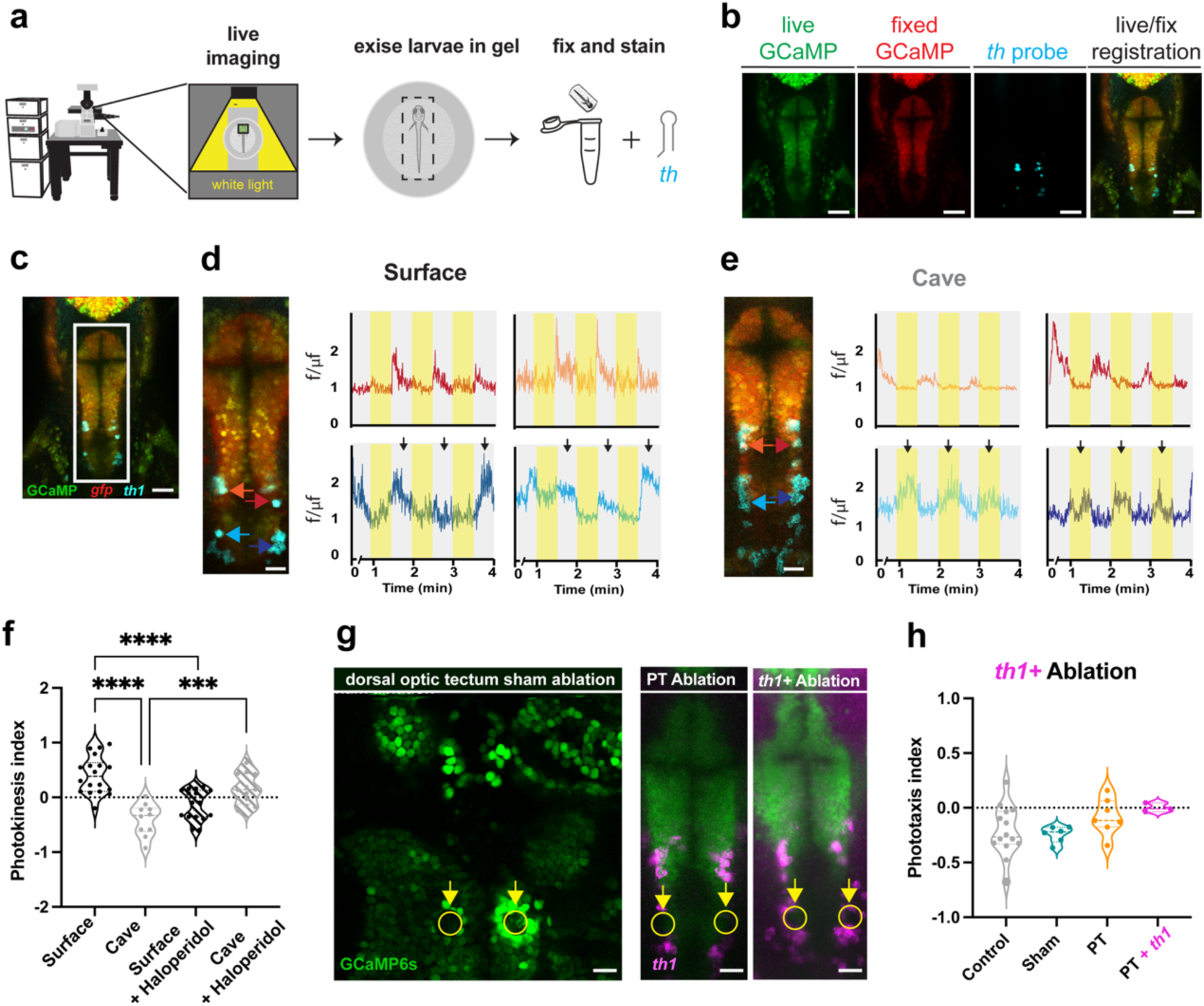
Light tuned PT neurons are dopaminergic and photokinesis is extinguished by dopamine antagonism or physical ablation. **a** pipeline for live to fix imaging; neural activity is recorded, larvae are fixed in agarose and stained with RNA probes. **b** Stained larvae are then imaged and registered to a maximum projection of the live stack. **c-e** Medial projection of the posterior tuberculum with colored dots and arrows corresponding to neural activity traces for surface fish (**d**) and cavefish (**e**). Images resolutions are (**b&c**) 512 x 512, zoom 1.2, and (**d&e**) 512 x 128, zoom 2.4. **f** Violin plot showing photokinesis indices for surface fish, cavefish, surface exposed to haloperidol and exposed to haloperidol. Sample sizes, surface fish (n=18), cavefish (n=10), surface fish + haloperidol (n=15) and cavefish + haloperidol (n=12). **g** Examples of 2-photon sham, posterior tuberculum (PT) and *tyrosine hydroxylase* (*th*) positive ablations. **h** Photokinesis indices following ablation experiments. Sample sizes, cavefish (n=14), cavefish sham ablations (n=6), cavefish PT ablations (n=7) and cavefish PT +*th* ablations (n=3). Scale bars, 512 x 512 = 50 μm and 512 x 128 = 20 μm. Statistical significance represents a Tukey multiple comparisons corrected one-way ANOVA. *** = p<0.001, **** = p<0.0001

### Functional role of A11 dopaminergic neurons in the caudal posterior tuberculum modulating light evoked locomotion

To functionally test if dopamine plays a role in photokinesis behavior, we exposed surface fish and cavefish to the typical antipsychotic dopaminergic inhibitor, haloperidol, and then tested larvae for photokinesis behavior. We found that haloperidol abolishes photokinesis behavior in both surface fish and cavefish (Figure5f, Supplemental Tables S21&22), further supporting the notion that dopaminergic neurons play a role in photokinesis behavior. While these data reveal that dopamine is necessary for light sensitivity, the wide-spread nature of DA cannot confirm the A11 descending diencephalic system as the necessary neurons. To confirm that the A11 neurons were indeed those responsible for light preference, we used a multiphoton laser to physically and specifically ablate cavefish neurons in the caudal region of the posterior tuberculum and then determined whether these specific neurons impact photokinesis behavior (Supplementary Figure S5). Experimental ablations were targeted to the caudal portion of the tuberculum, with sham ablations targeted to dorsal visual regions of the brain as a control (Figure 5g, Supplementary Figure S5, Supplementary Table S19). In contrast to control groups, ablations to the posterior tuberculum reduced photokinesis behavior, while ablation of *th* expressing neurons provided the largest change in behavior (Figure 5h). Taken together, our molecular maps, coupled with dopamine antagonism and physical ablations, suggests that dopaminergic neurons of the posterior tuberculum play a key role in behavioral variation across populations of *Astyanax*.

## Discussion

Neural circuit evolution underlies the functional variation necessary for behavioral adaptation in a new environment. Behavioral adaptation has long been linked to anatomical and physiological changes in the brain, yet little is known about how behavioral evolves due to the complexity and lack of functional tools needed to study these relationships. This study tackles this impediment with a system that couples an evolutionary non-model vertebrate with cellular resolution functional tools. Our discovery that cavefish have evolved a light photokinesis allowed us to ask what brain regions are impacted and which neuronal subgroups could contribute to behavioral variation. The fact that all previously studied eyed fish exhibit dark photokinesis and that only cavefish exhibit an inversion, suggests that the development of light photokinesis is reflective of evolving in a cave environment [37, 41]. This study also continues to expand our knowledge of the neurological basis of trait evolution in cavefish and more generally for sensory systems in vertebrates.

### Light photokinesis provides an adaptive behavior for avoiding light exposure and predation

Light photokinesis has developed convergently in cave fauna across the globe, suggesting that light photokinesis or photophobia is a cave adapted behavior. IN these ecosystems, animals are adapted to the light deprived nature of caves, and become less adapt at surviving in above ground territories. Past studies have found that cave adapted insects and fish exhibit agitation and increased activity when stimulated by light, and tend to retreat to dark conditions, behaviors termed photophobia or photokinesis [27, 55–61]. These studies suggest that light evoked activity is likely an adaptation for staying away from cave entrances and karst windows (sunken cave roofs), which could in turn place these organisms in an environment they are not conditioned for. Photokinesis would therefore reduce predator interactions or keep the cave adapted organisms from venturing towards surface environments [55, 56, 58–61]. This includes a study where light photokinesis was observed in cavefish that maintained an intact pineal, supporting our photokinesis brain mapping that shows increased activity for surface and cave morphs during photokinesis [62]. Therefore, our experiments provide functional knowledge to contribute to ethologically observed behavior, suggesting that forebrain circuits underlying photokinesis are evolving to functionally change light evoked hyperactivity.

### Surface to cave hybrid behavior suggests that photokinesis is genetically inherited

Surface to cave hybridization provides a functional genetic test to determine whether phenotypes are generically encoded [28, 63]. A trait is considered to be genetically encoded when a hybrid populations traits span the range of pure surface and pure cave traits. We observed a wide-range of photokinetic responses in surface fish to cavefish F_2_ hybrids, from surface-like dark activating to cave-like light activating, providing evidence that development of light or dark photokinesis is genetically encoded. [64, 65]. Cave light photokinesis could be a consequence of fixed genetic variants that impact sensorimotor integration during larval development. However, the fact that these fish have evolved in complete darkness, suggests that light photokinesis could be an indirect result of a fixed allele, such as a change in a sensory integrative hub, reflecting a fitness benefit for gustatory, aural or vibrational senses.

### Variation in function of an existing neuronal subgroup potentially explains inversion of stimulus cue in cavefish

While our research suggests that tubercular neurons play a role in photokinetic behavioral variation, we have yet to confirm whether this variation is due to a novel subset of neurons or a functional change in a group of neurons with shared ontology. Sensory circuits can be modified by several different mechanisms, including the loss or gain of a receptor, changes in accessory structures of a neuron and inclusion or exclusion of interneuron subgroups [3, 16, 66]. Photokinetic circuit function could be caused by evolution of a neo-functional cell type, where the regulatory machinery is reprogrammed to produce alternative functions in comparison to an ancestral cell type [67, 68]. Alternatively, a major shift in opsin wavelength tuning expression of varying opsins across populations could provide a receptive/integrative mechanism for producing light photokinesis [69–71]. However, we have yet to look at the genetic and developmental mechanisms driving variation in the posterior tuberculum, including genes-of-interest that could be causing this inversion. Finally, we present evidence that dopamine plays a functional role in cavefish photokinesis behavior, however we have not functionally tested the role of A11 dopaminergic neurons in surface fish dark photokinesis behavior. Future work looking at the development and function of these circuits are needed to clarify the evolutionary mechanism producing this variation in behavior.

### Dopamine is linked to evolution of the photokinetic variation in cavefish

Dopaminergic neurons play integral roles in sensorimotor integration and behavioral state changes [11, 53, 54, 72, 73]. We identified A11 dopaminergic neurons in cavefish that are light responsive in comparison to dark responsive in surface fish. A11 dopaminergic neurons have been shown to be highly conserved, with studies in fish, rodents and primates sharing the same developmental gene regulatory network [53, 54, 72]. Previous work has shown that A11 dopaminergic neurons are one of the primary sources of descending dopamine to the spinal cord in mammals, and that their activation leads to descending motor signals and locomotion [54, 72]. In zebrafish, researchers showed associations between neuronal activity in A11 dopaminergic neurons and several stimuli. This included subclasses that were reactive during spontaneous behavior, mechanosensory stimuli and visual stimuli [74]. Specifically, that dorsal medial dopaminergic neurons are active during spontaneous behavior and that visually stimulated tuberal hypothalamic neurons were reactive to a forward moving stimulus. The authors speculate that tuberal hypothalamic neurons may modulate reticulospinal neurons during swimming or during the optomotor response [74]. Our ablation results show that the posterior tuberculum and A11 dopaminergic neurons could provide a regional mechanism for inverting phototaxis, however more functional work is needed to clarify functional roles in surface and cave populations.

### Mutations causing increased levels of dopamine in cavefish provide a mechanism for photokinesis variation

Variation in monoamine metabolism may provide a mechanism for inverting the cavefishes photokinetic response. Monoamine metabolism is necessary to produce and regulate the neuromodulators dopamine, serotonin and noradrenaline [75, 76]. Previous work looking at the monoamine synthesis system in *Astyanax*, revealed that cavefish have a mutation in the *monoamine oxidase* (*mao*) gene that contributes to high levels of dopamine, serotonin and noradrenaline [77, 78]. Alternatively, a mutation in the ocular and cutaneous albinism 2 (oca2) gene that underlies a pigment synthesis pathway, may also lead to high levels of dopamine and serotonin [63, 79]. These changes in monoamine metabolism were hypothesized to play a role in the ‘behavioral syndrome’ that is observed in cavefish, including changes in sleep, stress and feeding [77]. We believe that variation in monoamine metabolism could play a role in light photokinesis. However, further genetic and developmental studies are needed to determine the true cause of light photokinesis in cavefish.

In conclusion, we find that cavefish provide a unique model for studying the evolution of sensory processing and sensorimotor behavior. Our work highlights the importance of neural circuit evolution for processing environmental stimuli and functional adapting to a novel environment. This work supports a light sensory processing role for dopaminergic neurons of the posterior tuberculum, while future work on sensory processing in the forebrain of cavefish will lead to a more comprehensive understanding of the mechanisms underlying the evolution of sensory systems.

## Methods

### Fish maintenance and husbandry

Zebrafish and Mexican tetras were housed in the Florida Atlantic Universities zebrafish and Mexican tetra core facilities. Larval fish were maintained at 28**°**C (zebrafish) and 23**°**C (Mexican tetra) in system-water and exposed to a 14:10 hour circadian light:dark cycle. Zebrafish and Mexican tetras were cared for in accordance with NIH guidelines and all experiments were approved by the Florida Atlantic University Institutional Care and Use Committee protocol #’s ###. Astyanax mexicanus surface fish lines used for this study include, surface fish Texas and surface fish Texas *Tg(_zfish_elavl3:H2B-GCaMP6s*). cavefish lines used for this study include, Pachón cavefish, Pachón *Tg(_zfish_elavl3:H2B-GCaMP6s*). GCaMP transgenic lines were produced in a previous study [31]. Surface fish were crossed to Pachón to generate F_1_ hybrids, while F_1_ hybrid offspring were incrossed to produce surface to cave F_2_ hybrids.

### Photokinetic behavioral experiments

Photokinesis experiments were performed using a custom designed behavior chamber equipped with diffuser light box, illuminated from below by white light LED. Videos were collected at 25 fps with 1280 x 960 resolution using a Basler acA1300-60gm camera fitted with a 12 mm Megapixel lens or a Basler acA1300-200um (Edmund Scientific Co., Barrington, NJ, #33978) fitted with a 16 mm C-series NIR-VIS lens (Edmund Scientific Co., #67714) and fitted with a UV-Visual cutoff filter (Edmund Scientific Co., #65796). All photokinesis experiments were run using an ANSI SBS compatible 96 well microtiter plate. Data was collected and analyzed using the EthoVision XT software version 11.5 and 14 (Noldus Inc., Leesburg, VA). Fish were acclimated to the observation chamber at 23 °C in the dark for at least 30 minutes. All behavioral experiments were recorded between 11am and 3pm, with 3-5 independent trials per condition.

### Statistics

Ethovision generated raw data was binned in 30 second intervals and sorted into light cycles using custom matlab scripts. Average lights on or off transition periods were then imported into PRISM Version 8.1.2 (GraphPad Software, Dotmatics, Boston, MA). Behavior was assessed for normality and then statistically assessed using t-tests for wildtype and one-way ANOVAs for hybrid comparisons. Tukey p-value corrections were applied following statistical significance for ANOVAs.

### Brain-wide phospho-ERK mapping

ERK phosphorylation methods were carried out using a previously published analysis for comparing ERK phosphorylation (mitogen-activated protein kinase (MAP)-mapping [43]. Larvae exposed to lights-on for cavefish and light-off for surface fish, were used as the experimental condition, with the larvae from the opposite light condition being selected as the control state. Z-stacks were imaged on a Nikon A1R multiphoton microscope, using a water immersion 25x, N.A. 1.1 objective.

### Automated segmentation and brain region measurements

To segment and measure subregions of the brain, we utilized a previously published analytical method [45] that registers each larval brain to the *Astyanax* brain atlas, segments the brain using advanced normalization tools (ANTs; [80]) and computes regional brain volume for all brain regions in the *Astyanax* atlas (cobraZ; [81]). Finally, a simple linear regression was applied to determine whether brain region volume correlated with photokinesis indices.

#### Genetically encoded calcium imaging

Surface fish and Pachón cavefish GCaMP6s stable transgenic embryos were screened for GFP expression and grown to 6 dpf. Larvae were incubated in 10 ng/μL bungarotoxin dissolved in system water for 15 minutes before being moved to a recovery bowl with 250 mL of system water. Following 5 minutes of system water recovery, larvae were imbedded in 2% low melting point agarose dissolved in system water and placed on the confocal stage to acclimate for 15 minutes. Following acclimation, larvae were imaged for 4 minutes, one minute of no light following laser initiation, followed by 30 second intervals of light-on and light-off.

### Live GCaMP light on and off response analysis

Time series stacks were analyzed using mesmerize, a live imaging analyses suite that utilizes the program CaiMan for stimulus tunning and neuronal clustering. Timeseries stacks and regions of interests saved in FIJI were imported, along with stimulus periods, then motion corrected and saved as a dataframe. Dataframes were run through the raw data normalization, stimulus tuning, linkage matrix and hierarchical clustering functions (max_cluster=5), to cluster neurons according to light and dark responsiveness. These clusters were then mapped using the plot dataframe function and saved as a png.

### Molecular mapping of fixed to live images

Larvae were screened and GCaMP was imaged as described previously [31]. Following imaging, larvae were euthanized in tricaine, excised in agarose and fixed in 4% formaldehyde overnight at 4 degrees. Agarose blocks were then washed 3x with 1x PBS and stained using a previously published HCR in situ hybridization protocol [45]. HCR probes against surface fish and cavefish *tyrosine hydroxylase* (*th*) mRNA were generated and used to probe the posterior tuberculum. Blocks of stained agarose embedded larvae were then mounted in an imaging well using a drop of 2% low-melting point agarose and imaged on a Nikon A1R confocal microscope with a 20x long-working distance 0.9 NA water dipping objective. Images were then registered to live imaged slices using a previously published method [45].

### Dopamine antagonist exposure

A previously published protocol was used to expose larval fish to the dopaminergic antagonist Haloperidol [82]. 5 dpf larval fish were exposed to 500 nM Haloperidol dissolved in 0.1% DMSO (experimental) or 0.1% DMSO (control) overnight. The following day larvae were assayed for photokinesis behavior and analyzed as previously stated in the behavioral and statistical methods sections.

### 2-photon physical ablation

Larvae were anesthetized using 25 mg/L tricaine, embedded in 2% agarose and then subject to 2-photon laser ablation. Larvae brains were ablated using a Zeiss 20x, 1.0 NA, W Plan-Apochromat, water immersion (cat# 421452-9880). The following laser settings were used to physically ablate 10 μm^3^ bilateral circles; wavelength of 880 nm, 80% power, 8.3 mW, laser engaged for 5 seconds, 20x water immersion objective. Following ablation, larvae were extracted from agarose and allowed to recover overnight. The following day larvae were assayed for photokinesis behavior and analyzed as previously stated in the behavioral and statistical methods sections.

## Data Sharing

All data, statistical tables and custom codes can be found at https://drive.google.com/drive/folders/1rTEAh8wxZnpKtDJJhW1j9GKCqYEa_w-N?usp=drive_link

## Supporting information

All Supplemental Data

## Acknowledgements

We would like to thank the administration and staff at the Jupiter Life Science Initiative in the Department of Biology at Florida Atlantic University, especially Peter Lewis and Arthur Loppatto for overseeing the health and care of the FAU *Astyanax* fish facility. We would also like to thank Dr. Julia Dallman for the use of her behavior rigs at the University of Miami as an initial screen for this behavioral study.

## SUPPLEMENTARY FIGURES

**Supplementary Figure S1.**
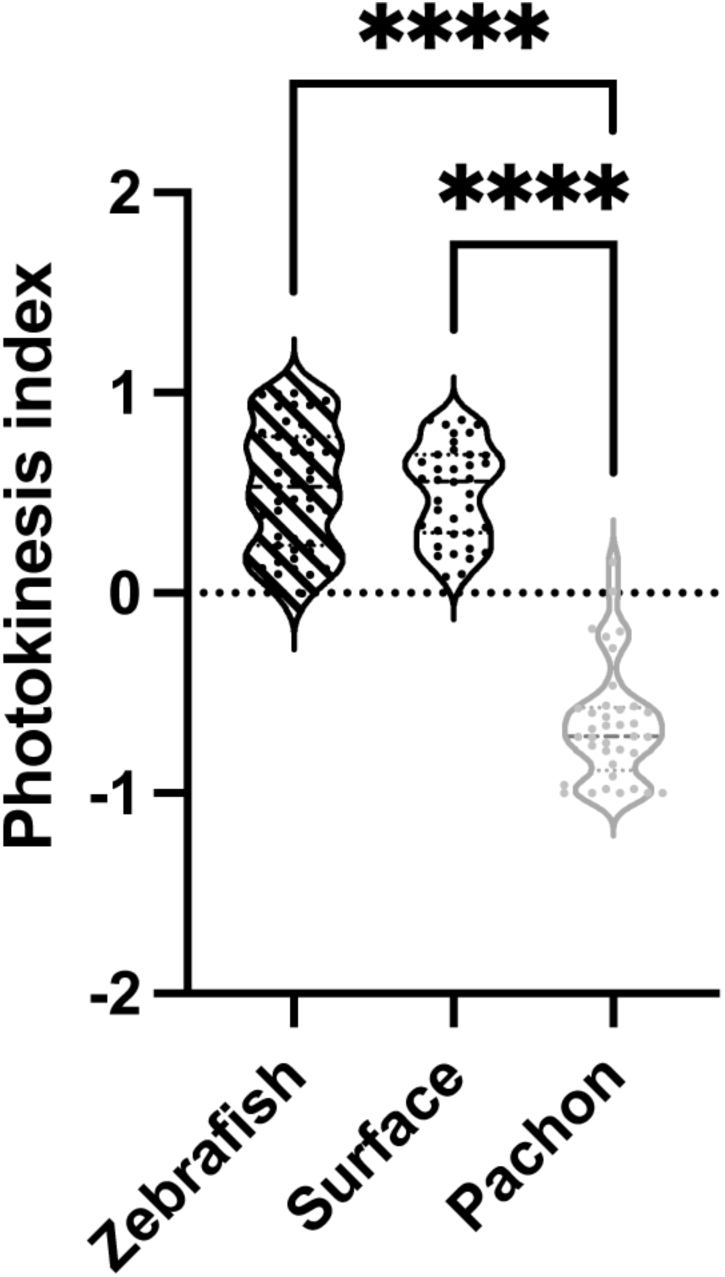
Zebrafish and surface fish exhibit positive photokinesis indices, while cavefish exhibit negative photokinesis indices. Sample size, zebrafish (n=47), surface fish (n=38), and cavefish (n=38). Statistical significance, **** = p<0.0001

**Supplementary Figure S2.**
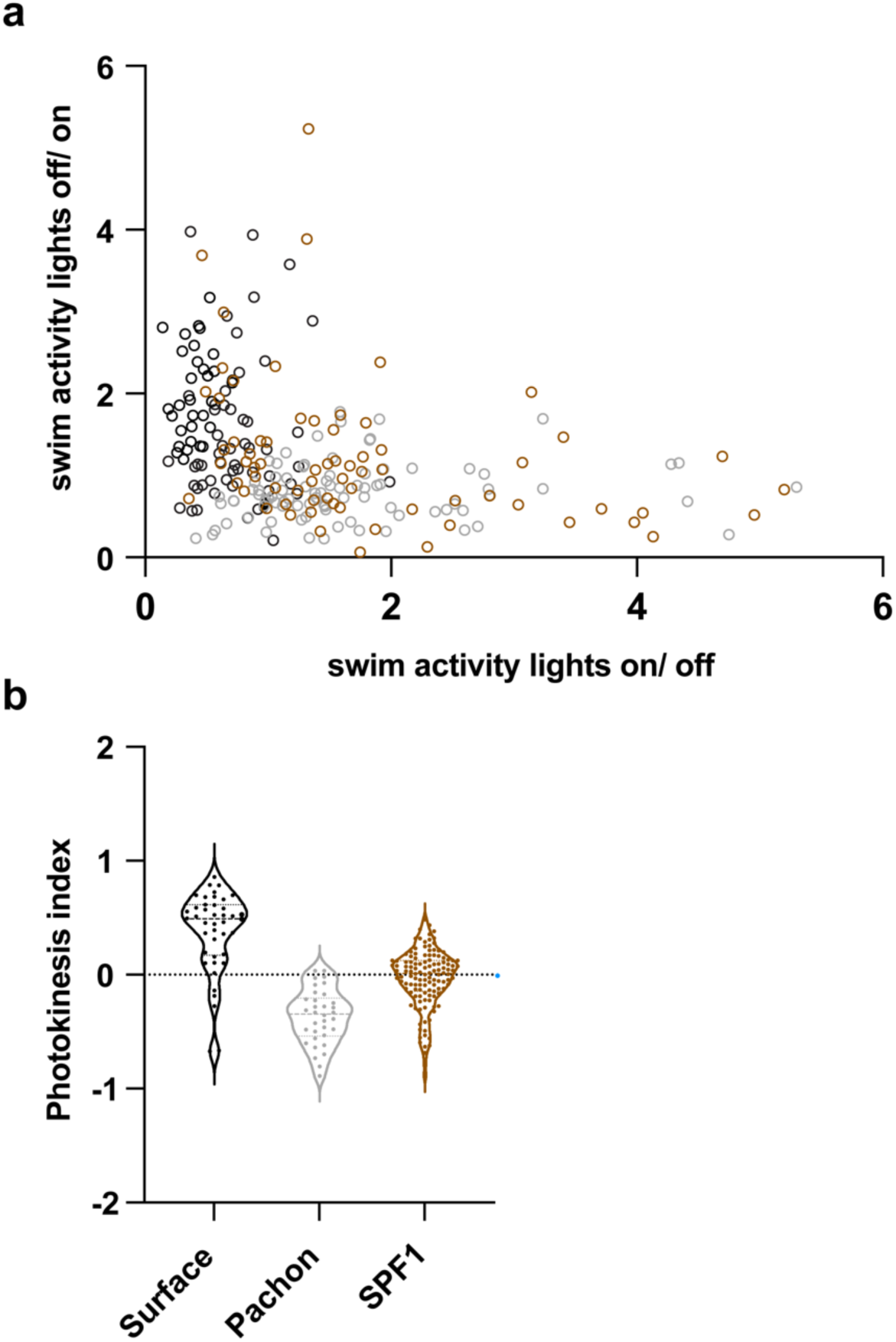
Surface to cave F_1_ hybrids exhibit a wide range of indices that suggests photokinesis is genetically inherited. **a** Scatter plot exhibiting the clustering of surface values along the y-axis, cavefish along the x-axis, and F_1_ hybrids spanning across both axes. **b** Box plots exhibiting positive photokinesis indices in surface fish, negative in cave and values spanning positive to negative in hybrids. Sample sizes, surface fish (n=42), cavefish (n=34), surface to cave F_1_ hybrids (n=126).

**Supplementary Figure S3.**
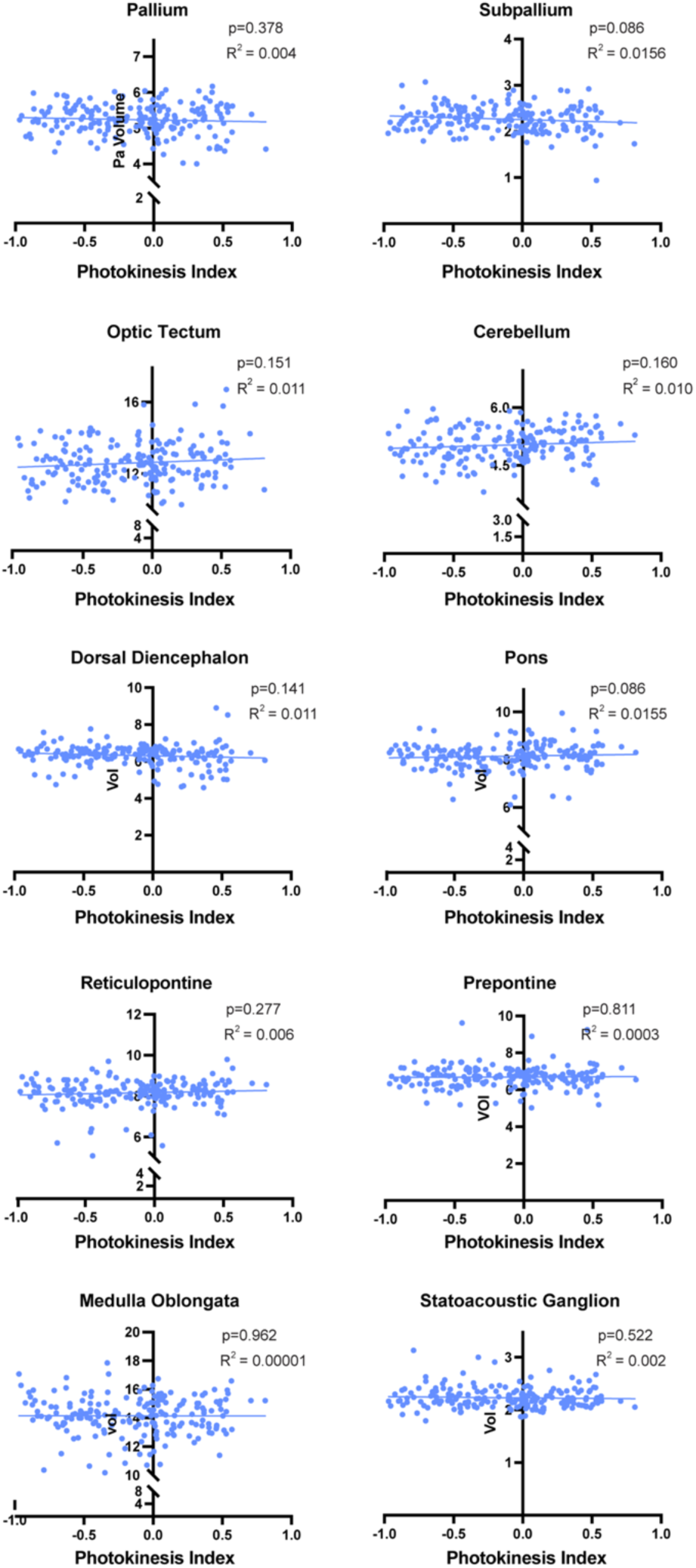
Brain regions exhibiting no correlation between brain region volume and behavior. X-Y scatter plots with linear regression, p values and R2 values. Sample size, surface to cave F_2_ hybrid n= 199.

**Supplementary Figure S4.**
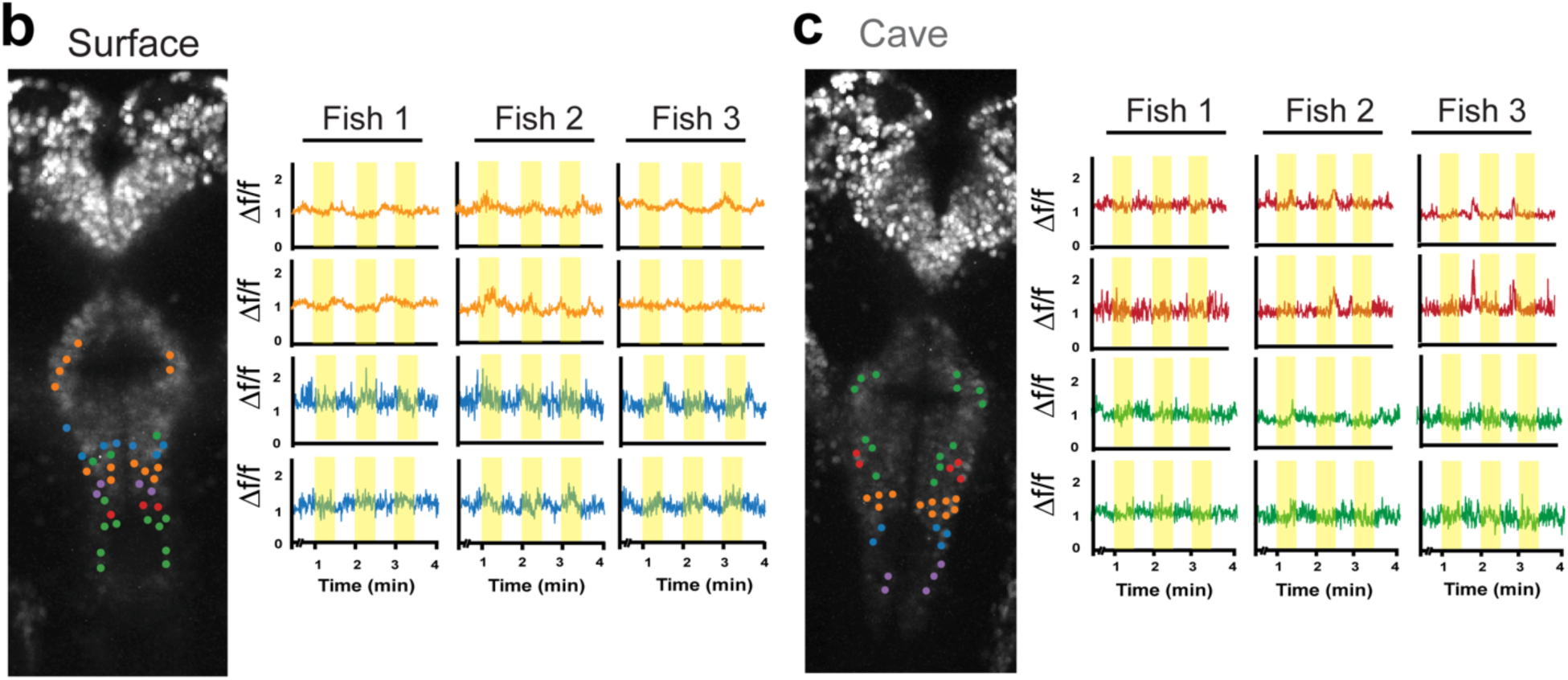
Rostral clusters of the posterior tuberculum do not exhibit light stimulus tuning for either lights-on or lights-off. Optical section of **a** surface and **b** cavefish posterior tuberculum to visualize neurons that are clustered in neural activity traces.

**Supplementary Figure S5.**
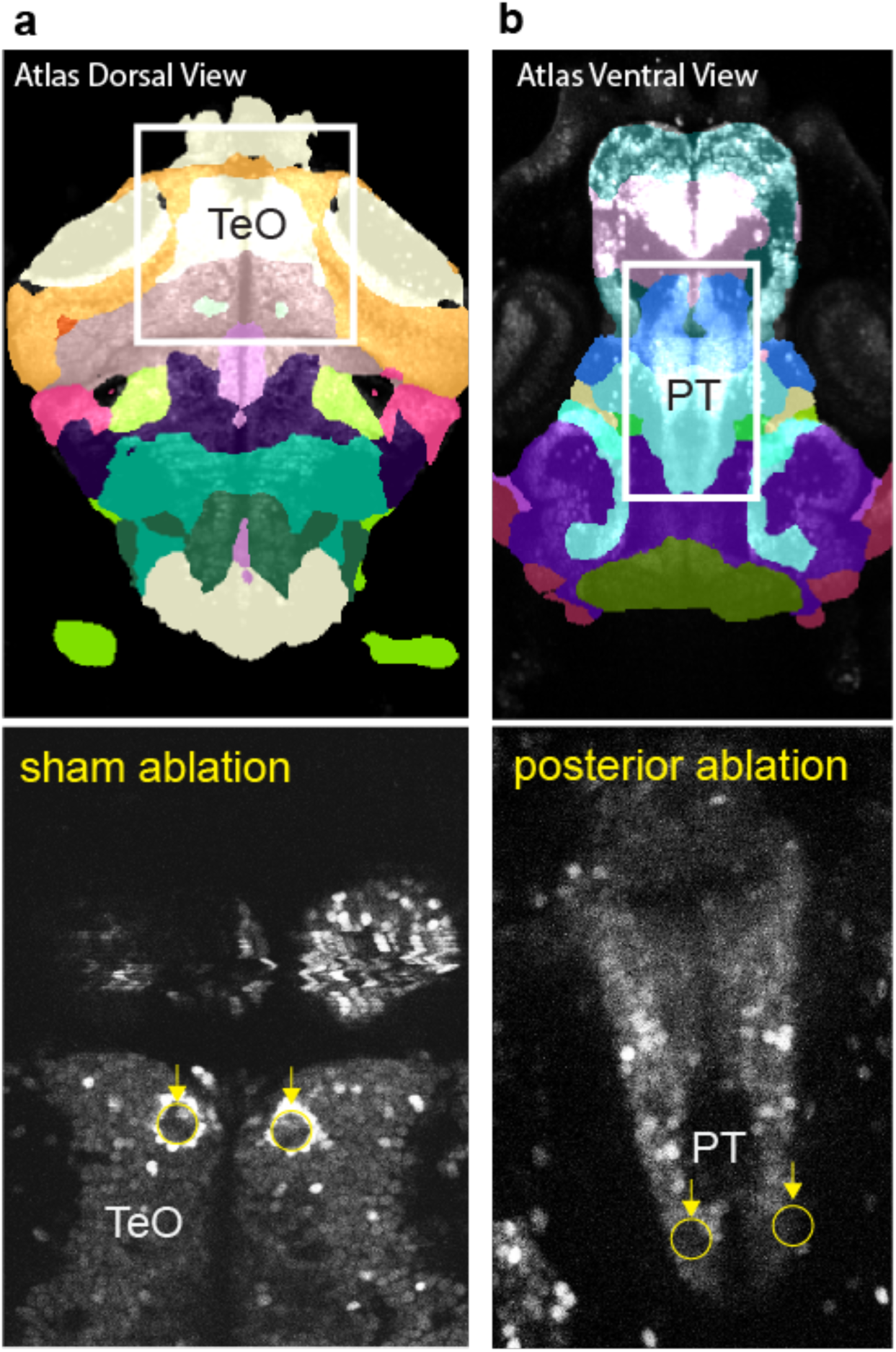
Examples of ablations in dorsal visual regions and the ventral posterior tuberculum. **a** Optical section of the *Astyanax* atlas to visualize the bilateral lobes of the optic tectum (TeO, above) and example of sham ablation (below) **b** Optical section of the Astyanax atlas to visualize the oval shaped posterior tuberculum (PT, above) and example of targeted ablation (below).

